# Impacts of pleiotropy and migration on repeated genetic adaptation

**DOI:** 10.1101/2021.09.13.459985

**Authors:** Paul Battlay, Sam Yeaman, Kathryn A. Hodgins

## Abstract

Observations of genetically repeated evolution (repeatability) in complex organisms are incongruent with the Fisher-Orr model, which implies that repeated use of the same gene should be rare when mutations are pleiotropic (i.e., affect multiple traits). When spatially divergent selection occurs in the presence of migration, mutations of large effect are more strongly favoured, and hence repeatability is more likely, but it is unclear whether this observation is limited by pleiotropy. Here, we explore this question using individual-based simulations of a two-patch model incorporating multiple quantitative traits governed by mutations with pleiotropic effects. We explore the relationship between fitness trade-offs and repeatability by varying the alignment between mutation effect and spatial variation in trait optima. While repeatability decreases with increasing trait dimensionality, trade-offs in mutation effects on traits do not strongly limit the contribution of a locus of large effect to repeated adaptation, particularly under increased migration. These results suggest that repeatability will be more pronounced for local rather than global adaptation. Whereas pleiotropy limits repeatability in a single-population model, when there is local adaptation with gene flow, repeatability can occur if some loci are able to produce alleles of large effect, even when there are pleiotropic trade-offs.

**Article summary:** Classical evolutionary theory predicts that genetically repeated evolution should be vanishingly rare in organisms where mutations affect multiple traits. In this article, we use simulations to study such pleiotropic mutations, and explore their effects on local adaptation in two patches under divergent selection. We find that migration between patches increases the likelihood of repeated evolution, even when there are fitness trade-offs imposed by pleiotropy.

## Introduction

Studies of adaptation commonly observe repeated genetic changes, where multiple species independently respond to a given selection pressure with mutations in orthologous genes (Chan *et al*. 2010; Bohutínská *et al*. 2021; Tittes *et al*. 2021). These patterns imply a lack of redundancy in the genes available for a selective response (Conte *et al*. 2012; Yeaman *et al*. 2018), and at first glance seem inconsistent with another common observation: that standing variation in quantitative traits is explained by a very large number of alleles of small effect (Fisher 1919; Boyle *et al*. 2017), which suggests a high level of redundancy in the genes contributing to quantitative traits. Is this seeming inconsistency a straw man, arising from the former pattern being driven by adaptation, with the latter pattern driven by mutation-selection balance? Understanding how evolutionary processes shape the kinds of mutations that contribute to adaptation remains a central question in biology.

In the early 20th century, theoretical work by R. A. Fisher demonstrated that the continuous phenotypic variation observed in populations could be explained by a large number of alleles inherited in a Mendelian manner (Fisher 1919), and that selection would favour small-effect changes at large numbers of loci (Fisher 1930). Genome-wide association studies in humans have provided empirical observations of standing variation congruent to Fisher’s models of adaptive trait architecture (reviewed in Visscher *et al*. 2017): Associations with hundreds or thousands of genetic variants explain only a modest proportion of trait heritability, with the remaining heritability attributable to even larger numbers of variants with effect sizes too small to detect with current cohorts (or possibly to rare variants that are excluded from many such analyses). But if variation in thousands of genes underpins a given trait, why would we ever observe orthologous genes contributing to adaptation in multiple species, when there are seemingly myriad ways to construct the same traits?

In his revisiting of Fisher’s model, Kimura (1983) demonstrated that although smaller effect mutations are more likely to be favourable, beneficial mutations of small effect are less likely to fix, as genetic drift biases the contribution of intermediate-effect loci to adaptation. Later, Orr (1998) showed that effect sizes of mutations fixed during an adaptive walk should be exponential, illustrating the importance of large-effect mutations early in bouts of adaptation to a new and distant environmental optimum. Perhaps the simplest explanation for repeated genetic adaptation is that alleles of large effect are disproportionately likely to contribute to early phases of adaptation (e.g. because of their fixation probabilities), but only a subset of loci are able to generate alleles of large effect (Orr 2005). Repeated gene use would then occur if there is long-term conservation of the genotype-phenotype map and the potential for particular loci to generate alleles of large effect. This kind of heterogeneity in mutational potential among loci is consistent with the omnigenic model (Boyle *et al*. 2017; Liu *et al*. 2019) draws on data from gene expression and per-locus estimates of heritability to distinguish between a relatively small number of ‘core’ genes of large effect and a much larger number of ‘peripheral’ genes of small effect. Certainly, large-effect QTL have been identified in both experimental evolution studies (e.g. McKenzie and Batterham 1994) and natural populations (e.g. Doebley 2004; Shapiro *et al*. 2004), and genomic footprints of selective sweeps (Maynard Smith and Haigh 1974; Kaplan *et al*. 1989) provide evidence for strong selection at individual loci (Barghi and Schlötterer 2020; Schluter *et al*. 2021). Among these examples, pesticide resistance in insects and pelvic reduction in sticklebacks display striking patterns of repeatability (Shapiro *et al*. 2004; Ffrench-Constant 2013). When there is spatially divergent selection, migration may further increase the likelihood of large-effect alleles contributing to adaptation and repeatability, as the contributions of small-effect alleles are disproportionately limited by the swamping effect of gene flow (Yeaman and Whitlock 2011). Consequently, repeatability in the genetic basis of local adaptation is expected to frequently involve large-effect mutations, particularly when gene flow is high or drift is strong, as long as these processes do not overwhelm selection (Yeaman *et al*. 2018). It should be noted that repeated adaptation tends to be particularly strong when bouts of adaptation occur from the same pool of standing variation (Ralph and Coop 2015; MacPherson and Nuismer 2017), but here we focus on independent bouts of adaptation without such shared variation.

While alleles of large effect may be favoured early in adaptation or when there is migration-selection balance, their contribution to adaptation can be limited by pleiotropy. In both Fisher’s (1930) and Orr’s (1998) models, mutations are modelled as vectors in multidimensional phenotypic space; therefore mutations with a large effect in a favourable dimension generally deviate too far from the optima in other dimensions, with serious fitness consequences (e.g. Boyle 1665). Chevin et al. (2010) expanded this model to incorporate distinct genes which could vary in their pleiotropic properties: specifically the number of traits that mutations would affect, and the correlation in effects of mutations on different traits (the latter being a property that can arise from organization of genes into networks; Hether and Hohenlohe 2014). They demonstrated that repeatability in the genetics of adaptation occurs in genes where negative fitness effects of pleiotropy are minimized.

Migration between populations experiencing divergent selection on a single trait can result in the homogenization of small-effect loci, whereas large-effect loci are more likely to be maintained as differentiated (Yeaman and Whitlock 2011; Tigano and Friesen 2016). Previous work on multitrait migration-selection models has focused either on contrasting the importance of linkage vs. pleiotropy (Chebib and Guillaume 2021) or used G-matrix approaches to explore adaptation at the quantitative genetic level (Guillaume 2011; MacPherson *et al*. 2015), so the effect of pleiotropy and migration-selection balance on repeatability at the gene level is unknown. While repeatability is expected to decrease with increasing trait dimensionality under a model of adaptation in a single population (i.e., without migration; Thompson *et al*. 2019), it is unclear how migration-selection balance will affect this, especially if genes differ in their effect size (as per Chevin *et al*. 2010) and if the optima of multiple traits under selection vary independently.

When selection acts on multiple traits, pleiotropy can have either a beneficial or deleterious effect on local adaptation, depending upon the relationship between how trait optima vary in space and how pleiotropic mutations tend to affect multiple traits. If there is a strong mutational correlation but only one trait is under spatially divergent selection and another is under spatially uniform selection, then the genetic correlation arising from mutations will tend to cause maladaptive divergence in the trait under uniform selection (Guillaume 2011). In this context, pleiotropy tends to have a deleterious effect on adaptation. By contrast, if multiple traits are all under divergent selection in the same direction as genetic correlations that arise from pleiotropic mutations, then pleiotropy tends to have a beneficial effect on adaptation. Any mismatch between mutational correlation and the spatial pattern of selection therefore poses an additional cost of pleiotropy beyond the costs represented in the Fisher-Orr model, so it is important to consider how this affects repeatability. It is unclear whether such trade-offs would effectively counteract the benefit for alleles of large effect observed in single-trait models of migration-selection balance.

Here, we use individual-based simulations of quantitative trait evolution examining how variation in pleiotropy and migration-selection balance interact to affect genetic repeatability. We build on previous models, which have considered adaptation in a single population following an environmental shift, by introducing a second population adapting to a divergent environment, and observing interactions between migration, effect size, and pleiotropy in bouts of local adaptation. We model different trade-offs in the fitness effects of mutations by varying the values of the trait optima in the two populations, so that divergence among the optima can run in parallel or in opposition to the direction of genetic correlations arising from mutation, allowing us to study scenarios where pleiotropy is beneficial or deleterious, respectively. To simplify the exploration of variation in mutation properties among genes (as per Chevin *et al*. 2010), we model two kinds of loci: a focal locus that can generate mutations of larger effect, and background loci that generate alleles of smaller effect. For these two classes of loci we separately vary the effect size and the mutational correlation (the relationship between effect sizes on different traits). This approach is not intended to closely model the architecture of a real trait, but rather allows us to focus on the relative contribution of two different classes of locus (e.g. core vs. peripheral genes under the omnigenic model; Boyle *et al*. 2017) and the effect of varying their characteristics independently. We predict that increasing effect size or decreasing deleterious pleiotropy (both the overall trait dimensionality as well as mutational correlation) at the focal QTL relative to the other QTL will increase repeatability. We also expect that increasing migration between demes will increase the repeatability observed, as large-effect alleles can maintain local adaptation in the face of gene flow (Yeaman and Whitlock 2011).

## Methods

To study the factors driving repeatability in independent bouts of adaptation, we performed Wright-Fisher, forward-time simulations in SLiM (v. 3.3.1; Haller and Messer 2019) with adaptation to an environment that varied across two patches connected by migration. Adaptation within each patch was driven by selection on two (or more) traits (e.g. *Z_1_, Z_2_*).

To gain insight into the parameters capable of driving repeatability at a particular locus and their interaction, we simulated a simplified genome: Traits could be affected by mutations at five genetically unlinked QTL; within QTL the probability of recombination occurring between one base and the next was 2.5×10^-7^. Properties were uniform across four QTL, while aberrant properties were assigned to a single ‘focal’ QTL, where parameter values could be varied independently of the non-focal QTL. For some parameters, simulations were repeated with a total of 20 QTL and one focal QTL. Each QTL consisted of 500 di-allelic loci with genotypic values in [0, a, 2a], at which mutations occurred at a rate of 1×10^-7^ per base pair per generation, resulting in an expected 10,000 mutations in each of two demes over the 20,000-generation simulation. This trait architecture is chosen to facilitate theoretical exploration of the relevant processes when there are two classes of genetic loci, rather than represent a biologically realistic trait.

QTL mutations affected one or more phenotypes (e.g. *Z_1_* and *Z_2_*); mutational effects for each QTL were drawn from a multivariate normal distribution with variance *σ^2^*, which determines the QTL effect magnitude (i.e., effect size), and covariance which was equal to the QTL mutational correlation multiplied by *σ^2^* (Fig. 1).

**Figure 1.**
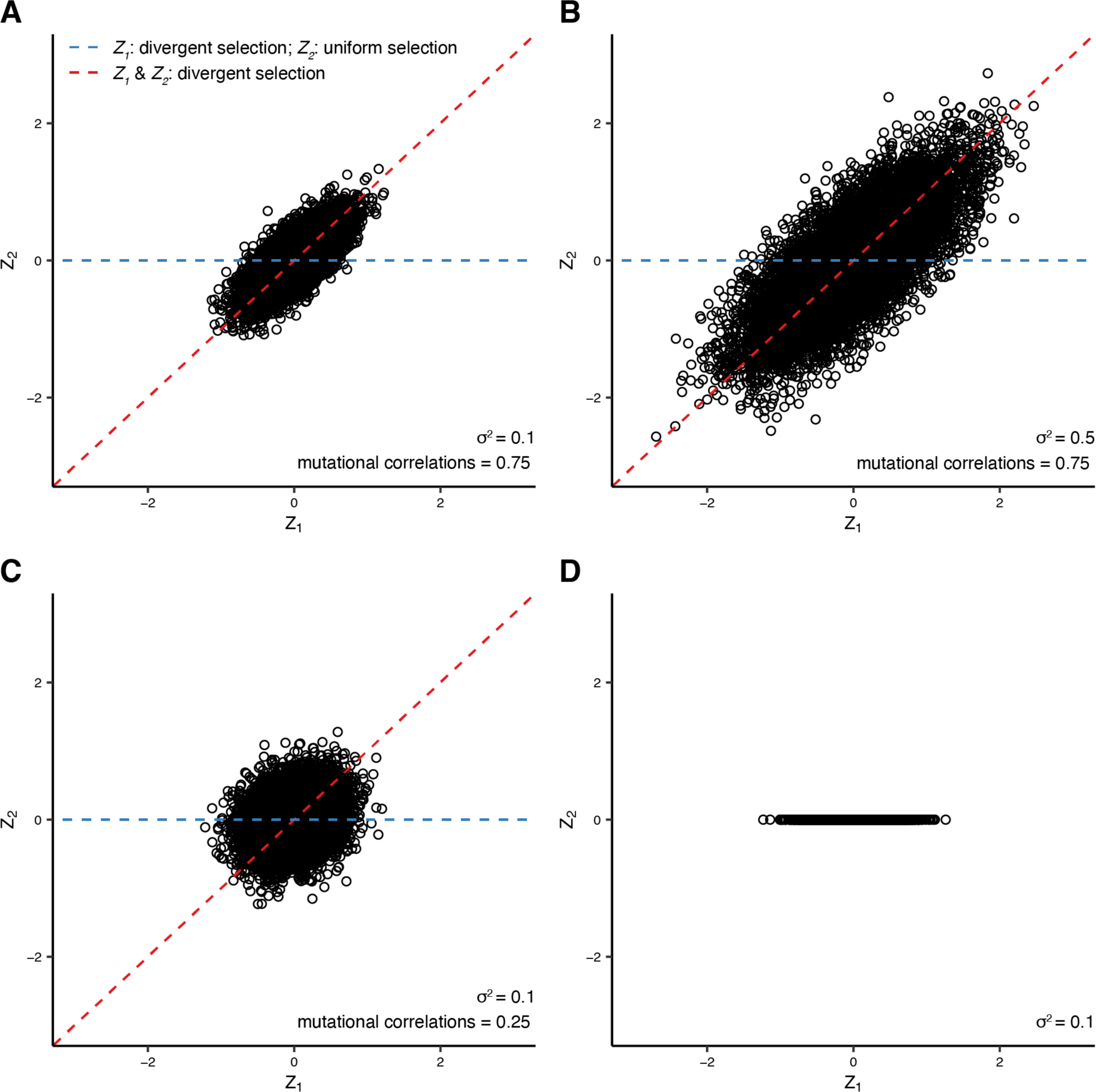
Different scenarios for patterns of selection on two traits (*Z_1_* and *Z_2_*), and correlation in mutation effects. Mutation effect sizes on traits *Z_1_* (x-axes) and *Z_2_* (y-axes) for 10,000 draws from distributions used to generate mutations. In A, *σ^2^* is 0.1 and the mutational correlation between traits is 0.75. In B, the mutational correlation is the same as A (0.75) but *σ^2^* is increased to 0.5. In C, *σ^2^* is the same as A (0.1), but the mutational correlation is relaxed to 0.25. In D, mutations have no effect on Z_2_. Red lines represent spatially divergent selection for both traits, along the same axis as mutational correlation (i.e. pleiotropy is beneficial), and blue lines represent spatially divergent selection for *Z_1_*, and spatially uniform selection for *Z_2_* (i.e., pleiotropy is deleterious). Red and blue pleiotropy guides are absent from panel D because mutations here only affect a single trait.

The following Gaussian function related individual fitness to phenotype *Z_i_*:

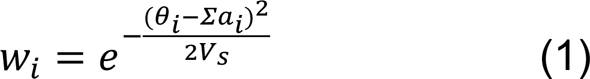

where *θ_i_* = the phenotypic optimum and *Σa_i_* = the sum across mutations of effect sizes on phenotype *Z_i_*, and *V_s_* = the variance in the fitness function, reflecting the strength of stabilizing selection (set at 125 for all simulations). Overall individual fitness was calculated as the product of *w_i_* across all phenotypes, so there was no correlational selection between pairs of phenotypes.

We simulated two demes (*d_1_* and *d_2_*), each composed of 1000 randomly-mating diploid hermaphroditic individuals. Phenotypic space was unitless and provided a relative scaling for the width of the fitness function and the magnitude of mutational effects. Both demes began the simulation with phenotypic optima of 0 for all phenotypes and ran with a burn-in for 20,000 generations. After the burn-in, phenotypic optima in *d_2_* were shifted and we tracked adaptive evolution over the following 20,000 generations. We focused on the case where in *d_2_*, the optimum for *Z_1_* was shifted to −10, while *Z_2_* (and optima for any other phenotypes) remained at 0 (Fig. 2C, D; top row). This configuration of trait optima means that when mutational effects on the two traits are correlated, mutations that are beneficial in one dimension will likely be deleterious in the other. For comparison, we also considered the case where all phenotypes in *d_2_* shift to −10, wherein pleiotropy generates mutations that are beneficial in all dimensions (Fig. 2A, B; top row). We varied the migration rate between *d_1_* and *d_2_* (0, 0.005, 0.05), *σ^2^* (0.1, 0.25, 0.5, 0.75, 1, 2, 3, 4, 5), mutational correlations (0, 0.25, 0.5, 0.75, 0.9, 0.99), and the number of phenotypes affected by the QTL (2, 5, 10).

**Figure 2.**
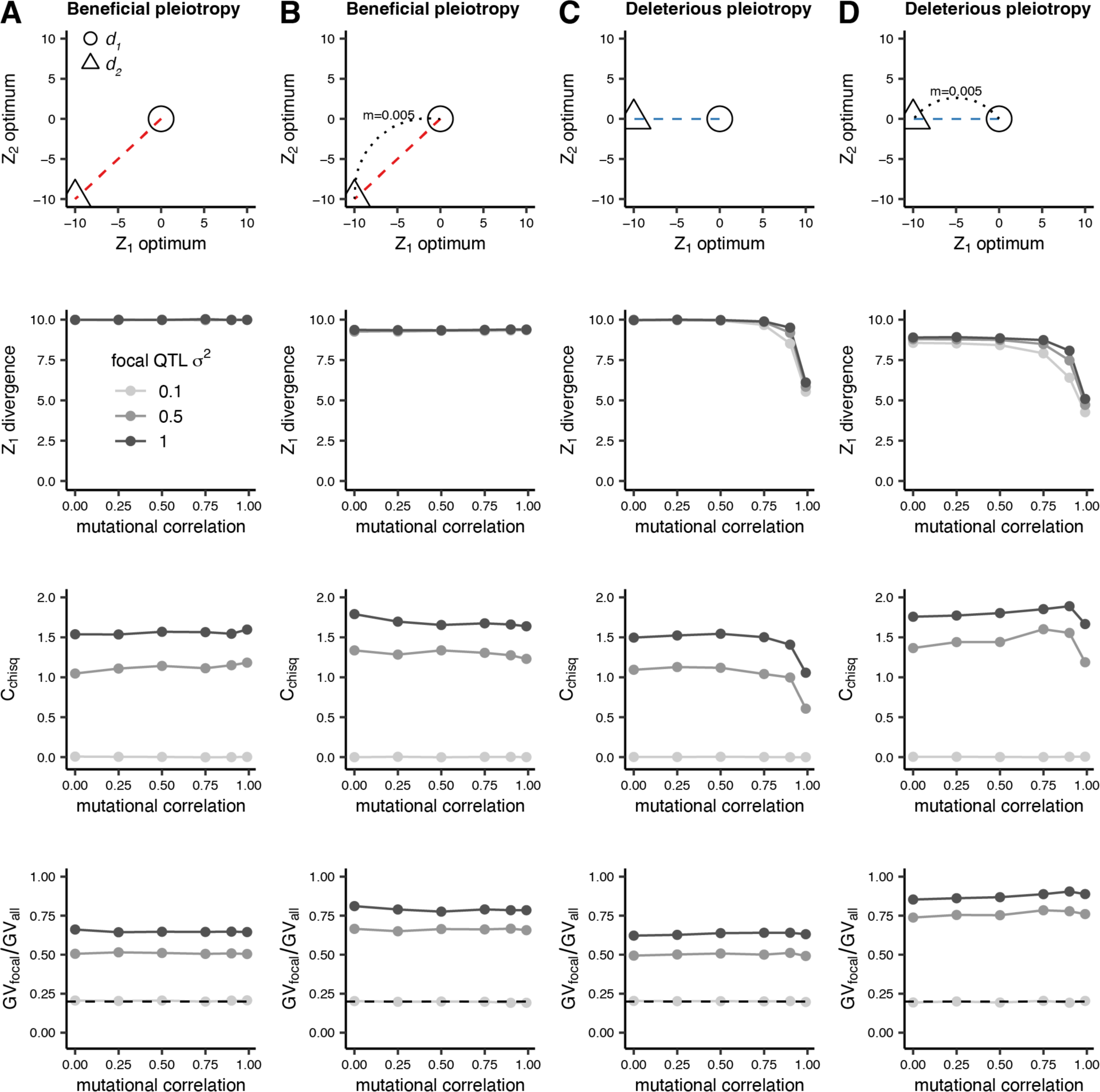
Divergence, repeatability (*C_chisq_*) and contribution of the focal locus (*GV_focal_ / GV_all_*) in *Z_1_* for high and low migration and beneficial and deleterious pleiotropy. The migration rates and pleiotropy are represented in the top row as circles [*d_1_*] and triangles [*d_2_*]. We model the situation where deme *d_2_* shifts to a new optimum in both traits *Z_1_* and *Z_2_* (divergent selection for both traits) without (A) and with (B) migration between *d_1_* and *d_2_*, and where *d_2_* shifts to new optimum in only trait *Z_1_* (divergent selection for one trait and uniform selection for the other) without (C) and with (D) migration between *d_1_* and *d_2_*. *σ^2^* at the focal QTL is varied, while *σ^2^* for non-focal QTL is 0.1. Mutational correlations are uniform at all QTL. Therefore when *σ^2^* = 0.1 focal and non-focal QTL have identical properties. The dotted line indicates *GV*_focal_ */ GV*_all_ = 0.2, the point at which the focal QTL explains the expected proportion of trait divergence under a null hypothesis and hence no repeatability is observed. Values greater than and less than 0.2 represent overuse and underuse of the focal QTL respectively (i.e., a greater than or less than expected contribution of the focal QTL to trait divergence). These simulations were run for 20,000 generations and 1000 replicates, with statistics calculated across replicates.

We investigated three main ways in which the types of mutation occurring at the focal QTL could be differentiated from those occurring at other loci:

1. A change in the QTL effect size, *σ^2^*, by altering the variance component of the variance-covariance matrix used to generate mutations (Fig. 1A c.f. B). This parameter was used to model a large-effect QTL at the focal QTL. *σ^2^* at non-focal QTL was always 0.1.
2. A change in mutational correlation by altering the covariance component of the variance-covariance matrix (Fig 1A c.f. C). This parameter models dependence between phenotypes and determines the probability that a mutation’s effect on one phenotype will have a corresponding effect on another.
3. A change in the number of phenotypes affected by a mutation by reducing the dimensionality of the variance-covariance matrix (Fig. 1A c.f. D). This models a situation where a QTL does not affect all of the phenotypes in the model.

To interpret the results of each parameter combination, we calculated the genetic value (*GV*) that a given genomic region (e.g., a QTL) contributes to phenotypic divergence using the formula:

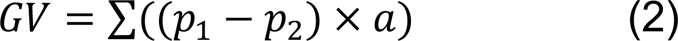

where *p_1_* and *p_2_* are the frequencies of a mutation in each deme, *a* is the size of the mutation’s effect on *Z_1_*, and values for each mutation are summed across a genomic region. To compare the relative impact of focal vs. non-focal QTL on divergence, we can then compare *GV* at these different classes of locus.

For each parameter combination we quantified the phenotypic divergence (the difference in mean phenotypes) between demes *d_1_* and *d_2_* at the divergently selected phenotype with 2×*GV_all_* (*GV* summed across all QTL). We also quantified disproportionate contributions of particular QTL to trait divergence (i.e., repeatability) across the 1000 simulation replicates. We used the *C_chisq_* statistic on QTL-specific *GV* with 1000 permutations (Yeaman *et al*. 2018), implemented in the dgconstraint R package (Yeaman *et al*. 2018) with the *pairwise_c_chisq()* function (i.e., each replicate is treated as an independent bout of evolution and compared in a pairwise fashion).

Briefly, χ^2^ was calculated across simulation replicates with:

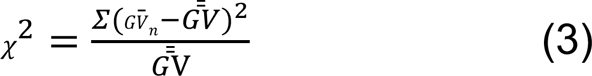

where 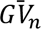 is the sum across simulation replicates of *GV* for the *n*^th^ QTL, and 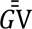 is the mean 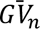 across all QTL.

The *C_chisq_* statistic was then calculated by using ×^2^ and 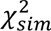 theresults of 1000 permutations of the data within each replicate:

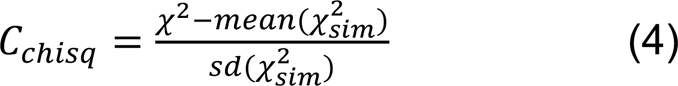

By this equation, when *C_chisq_*=0 we observe no more repeatability than would be expected by chance. The maximum value of *C_chisq_* varies with the number of QTL modelled: *C_chisq_*=2 for five QTL and *C_chisq_*≈4.36 for 20 QTL.

As an additional measure of the relative adaptive importance of the focal QTL, we calculated *GV_focal_ / GV_all_*, the proportion of *GV* summed across all QTL explained by *GV* summed across the focal QTL.

## Results

### Mismatch between the direction of mutational correlation and divergence in trait optima

We first present results for simulations where we expect pleiotropy to be beneficial (Fig. 2A, B) or deleterious (Fig. 2C, D), based on whether the multi-dimensional difference in trait optima between populations occurs along the same orientation as the mutational correlation (beneficial pleiotropy; red lines) or not (deleterious pleiotropy; blue lines). Repeatability of adaptation is uniformly low when all QTL have the same effect size (0.1; light grey points in Fig. 2), but increases substantially when the focal QTL has a larger effect size (darker points in Fig. 2).

When one population adapts to a new environment that differs in one phenotypic dimension but not the other, the axis of phenotypic divergence between the populations is not aligned with the axis of mutational correlation, and pleiotropy can be detrimental to adaptation (blue lines in Fig. 1; Fig. 2C, D). Based on Fisher’s model, we might expect large effect mutations at the focal QTL to be disfavoured in this scenario, as they would be increasingly likely to have maladaptive effects in one dimension. Surprisingly, we observe substantial repeatability regardless of the alignment between the correlation in mutation effects and divergence and trait optima (Fig. 2A vs. 2C; 2B vs. 2D), with only a slight reduction in repeatability for deleterious pleiotropy under the highest mutational correlations. The relative contribution of the focal locus to adaptive divergence (Fig. 2 bottom row) is also relatively insensitive to the strength of mutational correlation.

### The effect of migration

With both deleterious and beneficial pleiotropy, repeatability at the focal QTL tends to be higher with migration (Fig 2; Fig. S1), which again is particularly apparent when mutational correlations are strongest (Fig. 2C vs. 2D). This repeatability is driven by larger proportional contributions to divergence by the focal QTL, especially under increased migration (Fig. 2 bottom panels). As such, this increased repeatability with migration is likely due to the resistance to swamping of larger-effect mutations at the focal locus under migration-selection balance (as per Yeaman and Whitlock 2011). While deleterious pleiotropy with high mutational correlations impedes phenotypic divergence (Guillaume 2011), repeatability declines more in the absence of migration (compare high mutational correlation points in the *C_chisq_* plots between Fig. 2C and D). We observe repeatability in the presence of strong, deleterious pleiotropy, which occurs because high mutational correlations limit the rate of, but do not completely exclude the occurrence of mutations with fortuitous combinations of effects (Fig. S2A). Furthermore, there is a subtle effect across our simulations of the time point at which we sample: when *m*=0, repeatability decreases over time, whereas when *m*=0.005 it increases over time (Fig. S3). Thus, the observed effect of migration on repeatability would likely tend to increase over time.

### Variation among loci in mutational correlation

When pleiotropy is deleterious, reducing mutational correlation at the focal QTL relative to non-focal QTL may also allow it to more readily acquire adaptive mutations (thereby increasing repeatability). We modeled this by independently varying mutational correlations at the focal and non-focal QTL while holding the effect size of alleles constant (Fig. 3; Fig. S4). We observed increased repeatability when there were differences between mutational correlation values at focal and non-focal QTL, but no increase when all loci had the same correlation (Fig. 3). As expected, when the mutational correlation at the focal QTL was reduced relative to the non-focal QTL, repeatability of the focal QTL increased, and these patterns were consistent across migration rates (Fig. S4). The highest levels of repeatability were seen when the focal QTL had a relaxed mutational correlation against a background of high mutational correlation at non-focal QTL (i.e. 0.75 and particularly 0.9). This reflects the fact that mutational correlations need to be high to significantly limit the availability of mutations with fortuitous combinations of effects at the non-focal QTL and drive extreme repeatability at the focal QTL (Fig. S2B). When both focal and background QTL have the same correlation in their phenotypic effects there is no increase in repeatability, as shown in Fig. 3 where *C_chisq_* = 0 and where *GV*_focal_ */ GV*_all_ crosses the dashed line. When focal QTL mutational correlations are higher than the non-focal QTL, we also observe repeatability. In this case, however, repeatability is due to underuse of the focal QTL (i.e., it is contributing disproportionately less to divergence; Fig. 3B).

**Figure 3.**
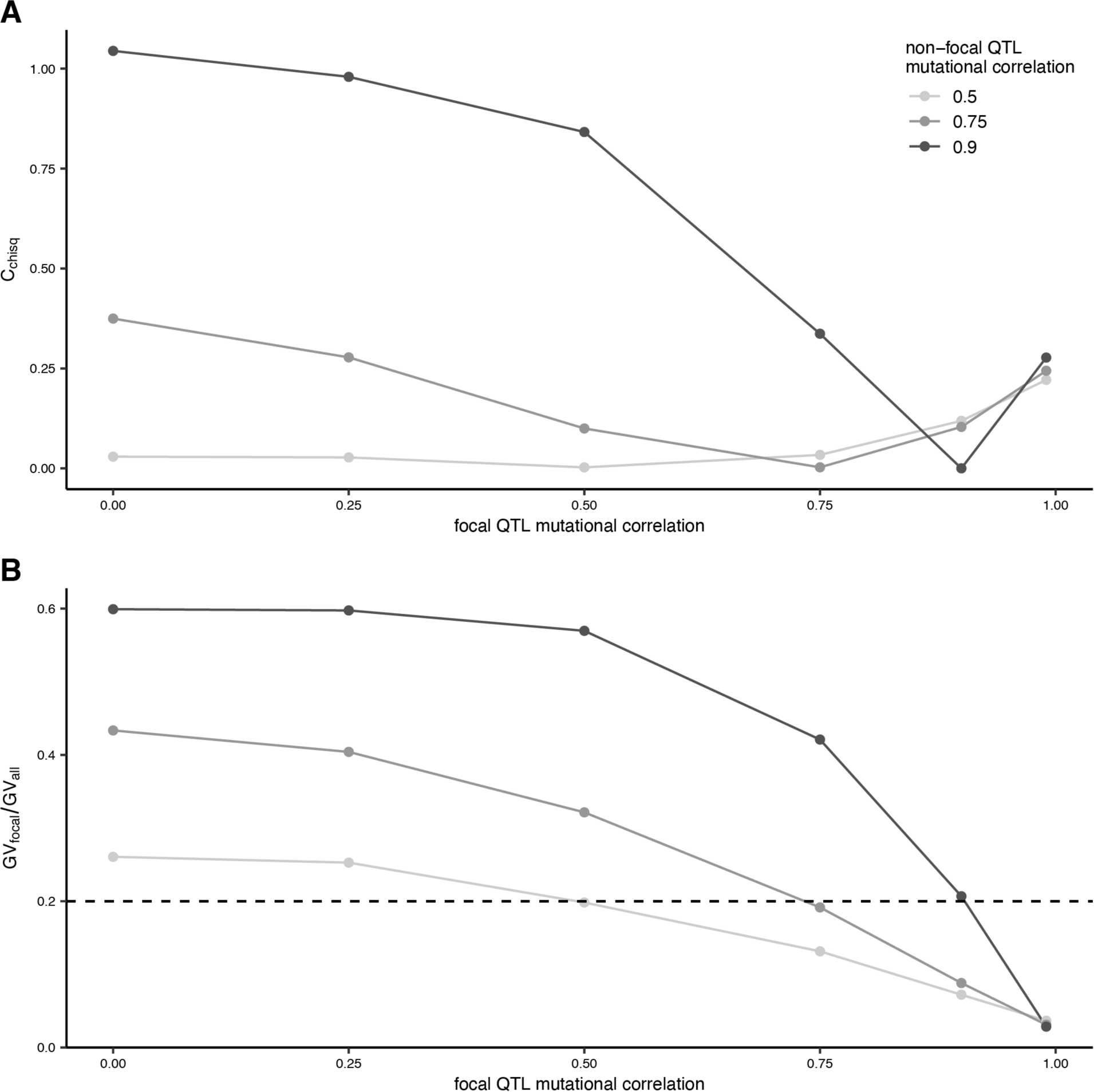
Repeatability (*C_chisq_*) in *Z_1_* against focal QTL mutational correlation for varying values of non-focal QTL mutation correlation (A), and the corresponding mean proportion of all *GV* explained by *GV* at the focal QTL in *Z_1_* (B). The dotted line indicates *GV*_focal_ */ GV*_all_ = 0.2, the point at which the focal QTL explains the expected proportion of trait divergence under a null hypothesis and hence no repeatability is observed. Values greater than and less than 0.2 represent overuse and underuse of the focal QTL respectively (i.e., a greater than or less than expected contribution of the focal QTL to trait divergence). These simulations use deleterious pleiotropy, a migration rate of 0.005, an *σ^2^*of 0.1 for both focal and non-focal QTL, and two phenotypes (one with divergent optima and the other with non-divergent optima), and were run for 20,000 generations and 1000 replicates, with statistics calculated across replicates.

### Variation in the number of trait dimensions

Up to this point our model of deleterious pleiotropy included only two phenotypes; one divergent and one non-divergent. To assess the robustness of these observations to an increase in the dimensionality of the phenotypes under selection, we increased the number of uniformly-selected phenotypes from one to nine. We first confirmed that migration increases repeatability when mutational correlations are absent (Fig. S5). We then reintroduced mutational correlations to the model and explored the effect of again varying the effect size (Fig. 4 middle row) and extent of pleiotropy (Fig. 4 bottom row) at the focal locus. When mutations can have pleiotropic effects in more dimensions, alleles of large effect are expected to experience greater constraints and contribute less to adaptation. Correspondingly, we observed that when pleiotropy was deleterious (i.e., there was a mismatch between the direction of mutation effects and the axis of divergence in phenotypic optima), we observed a decrease in repeatability with an increase in the number of traits affected by mutations but only when varying effect size at the focal locus (middle row Fig. 4C, D). However, similar to Fig. 2, the negative effect of pleiotropy on repeatability is less pronounced when migration is higher, presumably offset by the benefit for alleles of larger effect under migration-selection balance. When pleiotropy was beneficial, repeatability was not substantially affected by trait dimensionality (Fig. 4A, B).

**Figure 4.**
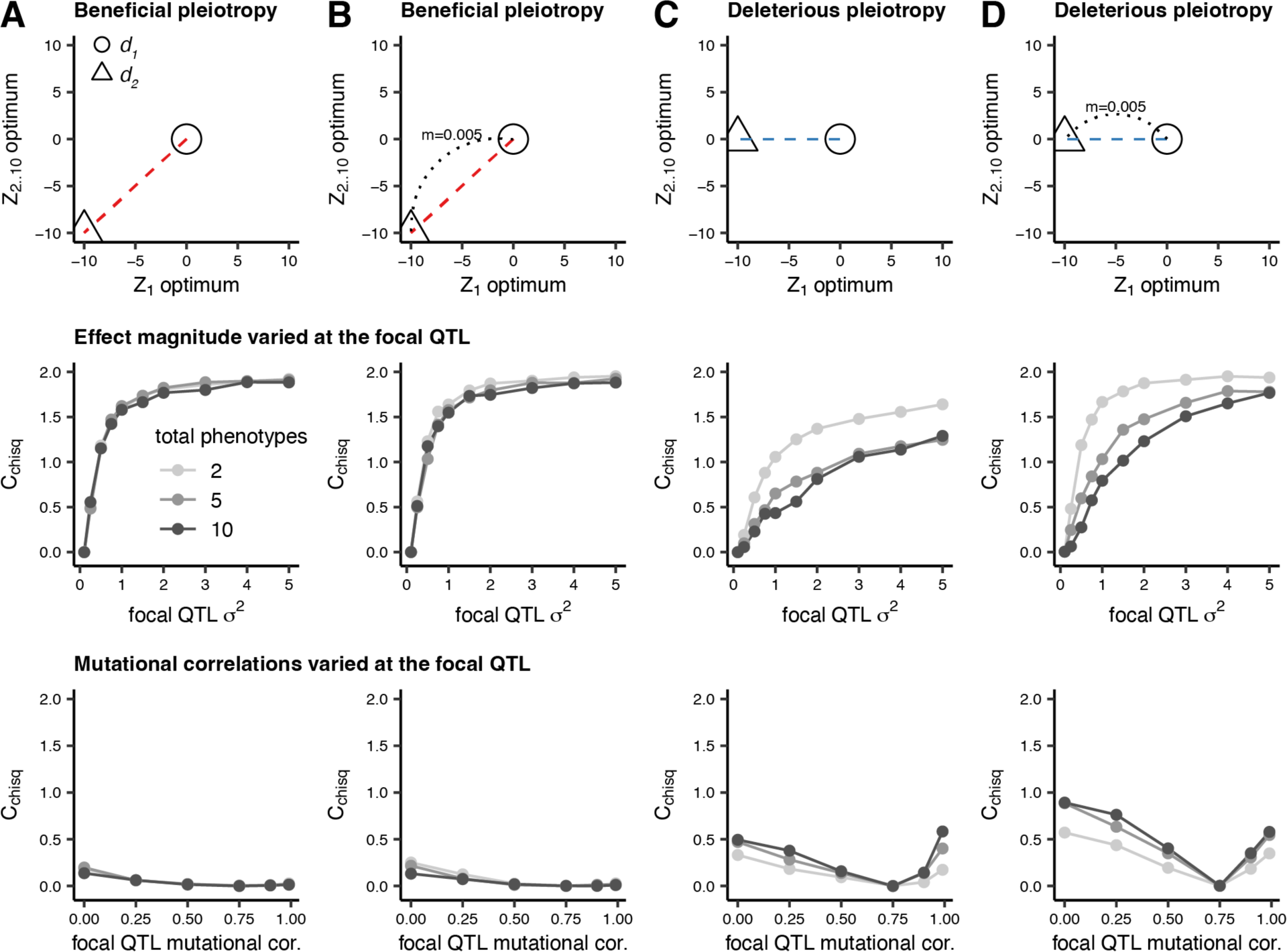
The effects of increasing trait dimensionality on repeatability (*C_chisq_*) arising from variation in mutational correlations or *σ^2^* at the focal QTL for four relationships between demes: where deme *d_2_* shifts to a new optimum in both traits *Z_1_* and *Z_2_* (a divergent selection for both traits) without (A) and with (B) migration between *d_1_* and *d_2_*, and where *d_2_* shifts to new optimum in only trait *Z_1_* (divergent selection for one trait and uniform selection for the other) without (C) and with (D) migration between *d_1_* and *d_2_*. Where *σ^2^* at the focal QTL is varied (middle row), *σ^2^* for non-focal QTL is 0.1 and mutational correlations at all QTL are 0.99. Where mutational correlations at the focal QTL are varied (bottom row), mutational correlations for non-focal QTL are 0.75 and *σ^2^* for all QTL 0.1. Simulations were run for 20,000 generations and 1000 replicates, with statistics calculated across replicates.

Finally, we investigated the case where mutations at the focal QTL affect fewer phenotypes than the non-focal QTL. In the two-phenotype model, this meant focal QTL mutations would only affect the divergent phenotype; in the five and ten-phenotype models, focal QTL mutations affected the divergent phenotype and one fewer non-divergent phenotypes than non-focal QTL (Fig. 5). Interestingly, with this difference in breadth of QTL effect between focal and non-focal QTL, the effect of trait dimensionality on repeatability is reversed, with the greatest repeatability seen at the highest number of trait dimensions. This is because pleiotropy at higher trait dimensions is more limiting, and hence relaxation of pleiotropy more beneficial. As before, with high mutational correlation between phenotypic effects, high levels of repeatability at the focal QTL are observed, however when mutational correlations are weak or absent, very little repeatability is observed.

**Figure 5.**
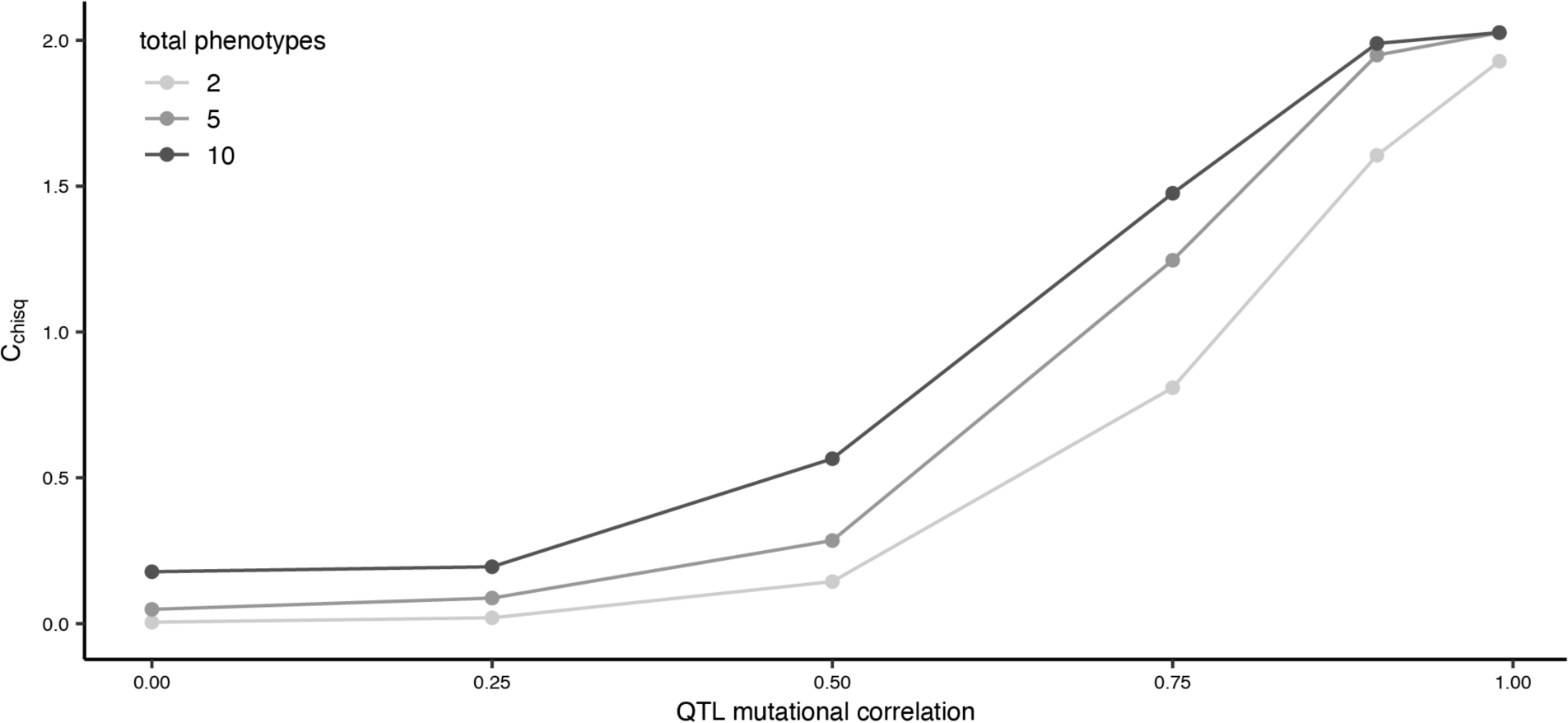
The effect of mutational correlation on repeatability when all QTL share the same mutational correlation and effect size, but the focal QTL affects the divergent phenotype (*Z_1_*) and one fewer non-divergent phenotypes than the non-focal QTL. Shades indicate the total number of phenotypes in the simulation (two with one non-divergent phenotype, five with four non-divergent phenotypes and ten with nine non-divergent phenotypes). These simulations use deleterious pleiotropy, a migration rate of 0.005, an *σ^2^* of 0.1, and were run for 20,000 generations and 1000 replicates, with statistics calculated across replicates.

## Discussion

Our simulations provide important insights for studies of local adaptation. Firstly, in the presence of adequate levels of migration, repeatability is expected to occur across lineages undergoing local adaptation to similar optima, even if strong pleiotropic relationships oppose the direction of divergence in phenotypic space (Fig. 2). Secondly, repeated use of a QTL in multiple lineages may arise if the QTL has a disproportionately large effect size (Fig. 2), or if deleterious pleiotropy at the QTL (either the amount of correlation between traits or the number of traits affected) is relaxed (Fig. 3, 5). Finally, for mutational correlations between divergent and non-divergent traits to influence repeatability, the correlations must be very high, so that fortuitous pleiotropy-breaking mutations are substantially limited (Fig. S2).

If all loci have equal pleiotropy and equal effect size, repeatability is not expected (light grey points in Fig. 2). Thus, the observation of repeated evolution (reviewed in Conte *et al*. 2012) clearly rejects a simple model where the effects of all loci are equal and interchangeable. While this may seem a “straw-man” model, it helps provide context to interpret theory built on the infinitesimal assumption. Such theories make an unrealistic assumption (that all loci have small and interchangeable effects) in order to generate testable predictions. Models based on the infinitesimal assumption should not be taken to imply that all loci necessarily have equal and vanishingly-small effects, but just that such an assumption allows for tractable analysis. It is therefore critically important to situate the results of this powerful family of models within the problematic observation that some loci clearly have large and repeatable contributions to adaptation. It will be interesting to continue to harmonize results from empirical data, classic population genetic models, and the powerful approximations that can be derived using the infinitesimal assumption (Turelli and Barton 1990; Barton *et al*. 2017; Hayward and Sella 2022).

To explore the effect of variation in pleiotropy on repeatability, we varied the properties underlying the mutations generated at focal vs. non-focal QTL. Firstly, we demonstrated that an increase in effect magnitude (*σ^2^*) of a QTL will produce patterns of repeatability, which is consistent with previous theoretical (Chevin *et al*. 2010) and empirical observations (e.g., Schlenke and Begun 2004; Rosenblum *et al*. 2004). Both mutational correlations and migration can force adaptation away from phenotypic optima along ‘genetic lines of least resistance’ (Schluter 1996; Guillaume 2011). Correspondingly, when the direction of mutational correlation is counter to the direction of divergence in optima, we see a reduction in trait divergence between demes as mutational correlation is increased, and a reduction in repeatability, which is less pronounced under higher migration, due to the benefit for alleles of large effect (Fig. 2). Similar to previous findings Thompson *et al*. (2019), we observe a reduction in repeatability with increased number of traits (Fig. 4), consistent with the expectation that when mutations can have pleiotropic effects in more dimensions, alleles of large effect experience greater constraints and contribute less to adaptation. However, if some loci affect fewer traits than others, we find the opposite pattern with higher repeatability at increasing trait dimensionality (Fig. 5). This is because when trait dimensionality is increased, pleiotropy is more limiting, and hence relaxation of pleiotropy more beneficial. Congruent with the findings of Chevin *et al*. (2010) who examined single populations, we also found that repeatability is increased if some loci have reduced mutational correlations (Fig. 3).

By simulating two demes linked by varying amounts of migration, our models address a common situation in local adaptation: Individuals in one population may experience local environmental shifts; they must therefore adapt to new optima for some phenotypes, while retaining existing optima at others. Previous research (Griswold 2006; Yeaman and Whitlock 2011) demonstrated that migration concentrates the genetic architecture of local adaptation and favours alleles of larger effect. Correspondingly, we find that high migration rates favour adaptation driven by mutations at a locus of larger effect when beneficial pleiotropy is present (bottom row of Fig. 2A c.f. B) and find that this still holds when pleiotropy is deleterious (bottom row of Fig. 2C c.f. D; Fig. S1). Thus, the tendency for mutations to have deleterious pleiotropy does not necessarily limit the tendency for loci of large effect to drive adaptation under migration-selection balance. Rather, adaptation tends to be achieved by alleles that happen to have less deleterious effects on the trait under spatially uniform selection (Fig. S2), which occur less often under strong mutational correlations. We also find that migration increases repeatability arising from variation in pleiotropy among focal vs. non focal QTL (Fig. S4). This is because repeatability is driven by the net effect of selection on a QTL. Under migration-selection balance those QTL with larger net beneficial effects (i.e., weaker deleterious pleiotropy) will be maintained as differentiated unless migration is so high that swamping occurs. Thus our results suggest that repeatability will be more pronounced in cases of local rather than global adaptation.

Guillaume (2011) utilized a similar two-patch design to investigate the effects of pleiotropy and migration on population divergence of phenotypes. He demonstrated that combinations of migration and pleiotropy can drive divergence between demes at phenotypes that share the same optima in both demes, as long as the phenotypes are sufficiently correlated with other divergently selected phenotypes. We observe similar patterns in our simulations: increasing levels of mutational correlations and migration reduce differentiation between demes at the divergent phenotype, and increase differentiation between demes in phenotypes not under divergent selection (Fig. S4). Perhaps most surprisingly, we show that this reduced phenotypic differentiation does not necessarily limit genetic repeatability, as high *C_chisq_* values are observed in simulations where pleiotropy and migration have substantially limited the divergence between demes (Fig. 2, Fig. S4). It seems that the mutational correlations, despite appearing quite high, do not actually pose substantial constraints to the evolution of genetic architecture, as they basically reduce the effective mutation rate in particular directions, but do not completely prevent such mutations from occurring.

Our simulations make a number of assumptions that are almost certainly violated in natural populations exhibiting repeatability. Firstly, we treat each simulation replicate as if it were a different species representing an independent bout of adaptation, and we assume complete orthology between QTL in replicates and that orthologous QTL retain corresponding effect magnitude and pleiotropic properties. In nature, divergence between species limits studies of repeatability to the orthologous portions of their genomes, and the effects of adaptation in non-orthologous regions has not been addressed here. Secondly, we have simulated both the initial phenotypic optima (to which both demes start our simulations adapted) and the divergent phenotypic optima as identical between replicates. Populations adapting to similar environments will not share identical phenotypic optima, which is important for the interpretation of our results, as Thompson *et al*. (2019) observed that repeatability declines rapidly as the angle between phenotypic optima increases, a pattern that increases with trait dimensionality. Furthermore variation between QTL in mutation rate, retention of standing variation and patterns of linkage disequilibrium may all affect the likelihood of repeatability, but we have held these parameters constant in our simulations. Finally, it is unclear what values mutational correlations may take in natural systems. In the absence of empirical estimates of mutational correlations, empirical estimates of genetic correlation between 0.3 and 0.8 (Maloy and Hughes 2013; Gidziela *et al*. 2023) suggest that the more extreme mutational correlations simulated in this work may not commonly occur in nature.

The simulations presented here also use a simplified genome architecture: four QTL with uniform properties and a single QTL with aberrant properties, and between two and ten traits. This system pales in comparison to the thousands of genes (exhibiting near-global pleiotropy) which contribute to traits under the omnigenic model (Boyle *et al*. 2017; Liu *et al*. 2019). Contrastingly, a meta-analysis of gene knockout experiments in *Saccharomyces cerevisiae*, *Caenorhabditis elegans* and *Mus musculus* (Wang *et al*. 2010) estimated pleiotropy to be far less pervasive: a median gene affects only one to nine percent of traits. Wang *et al*. (2010) also detected significant signals of modular pleiotropy (where subsets of genes affect subsets of traits), which would serve to simplify the architecture available for repeated adaptation. Simple genetic architecture enhances repeatability at a genome-wide level, and this study suggests that an even more modular architecture at some QTL will act to further magnify repeatability. While the nature of pleiotropic, quantitative traits in higher organisms remains unresolved, we expect our simple model to be applicable to more complex architectures (Yeaman *et al*. 2018), and repeating our simulations on models with 20 QTL yields comparable results (Fig. S6). Traits in real organisms will often be much more polygenic, with considerable variation among genes in the extent and direction of pleiotropy in their mutations, as well as the average effect size. If such differences in mutational characteristics of genes are conserved across deeper evolutionary time, then variation in their pleiotropic effects will strongly affect their contribution to repeatability.

This study demonstrates that in the absence of migration, pleiotropy limits repeatability, but when there is migration between environmentally divergent populations, repeatability can occur at large-effect loci. Our results are consistent with a pair of recent empirical studies examining the genes driving repeated global vs. local adaptation across multiple plant species. Genes driving local adaptation tended to be enriched for high pleiotropy (Whiting *et al*. 2023), while those driving global adaptation tended to be enriched for low pleiotropy (Nocchi *et al*. 2024).

This suggests that the interaction between pleiotropy and the tension between migration and selection will indeed impact the genetic architecture of adaptation. It will be interesting to see if other future studies find similar patterns.

## Data Accessibility Statement

Simulation Eidos scripts and analysis R code are available at https://github.com/pbattlay/pleio-sims/

## Acknowledgements

We thank Tim Connallon and Jacqueline Sztepanacz for valuable feedback on the manuscript.

## Funding

Funding was provided by a Human Frontier Science Program grant to KAH.

---

**Figure S1.**
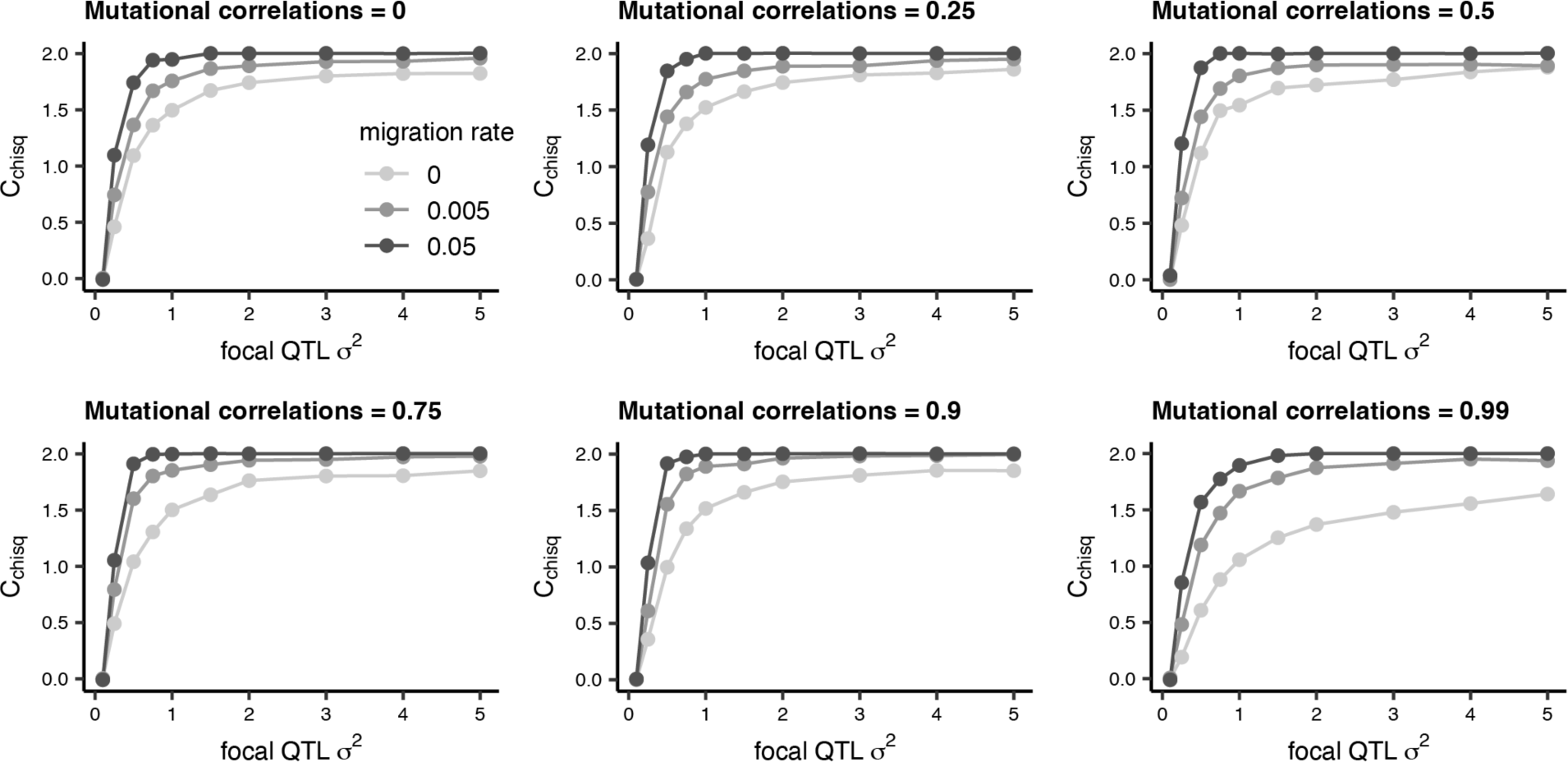
Repeatability (*C_chisq_*) in *Z_1_* against focal QTL *σ^2^* where the *σ^2^* for non-focal QTL is 0.1. These simulations use deleterious pleiotropy, two phenotypes (one divergent and one non-divergent). Simulations were run for 20,000 generations and 1000 replicates, with statistics calculated across replicates.

**Figure S2.**
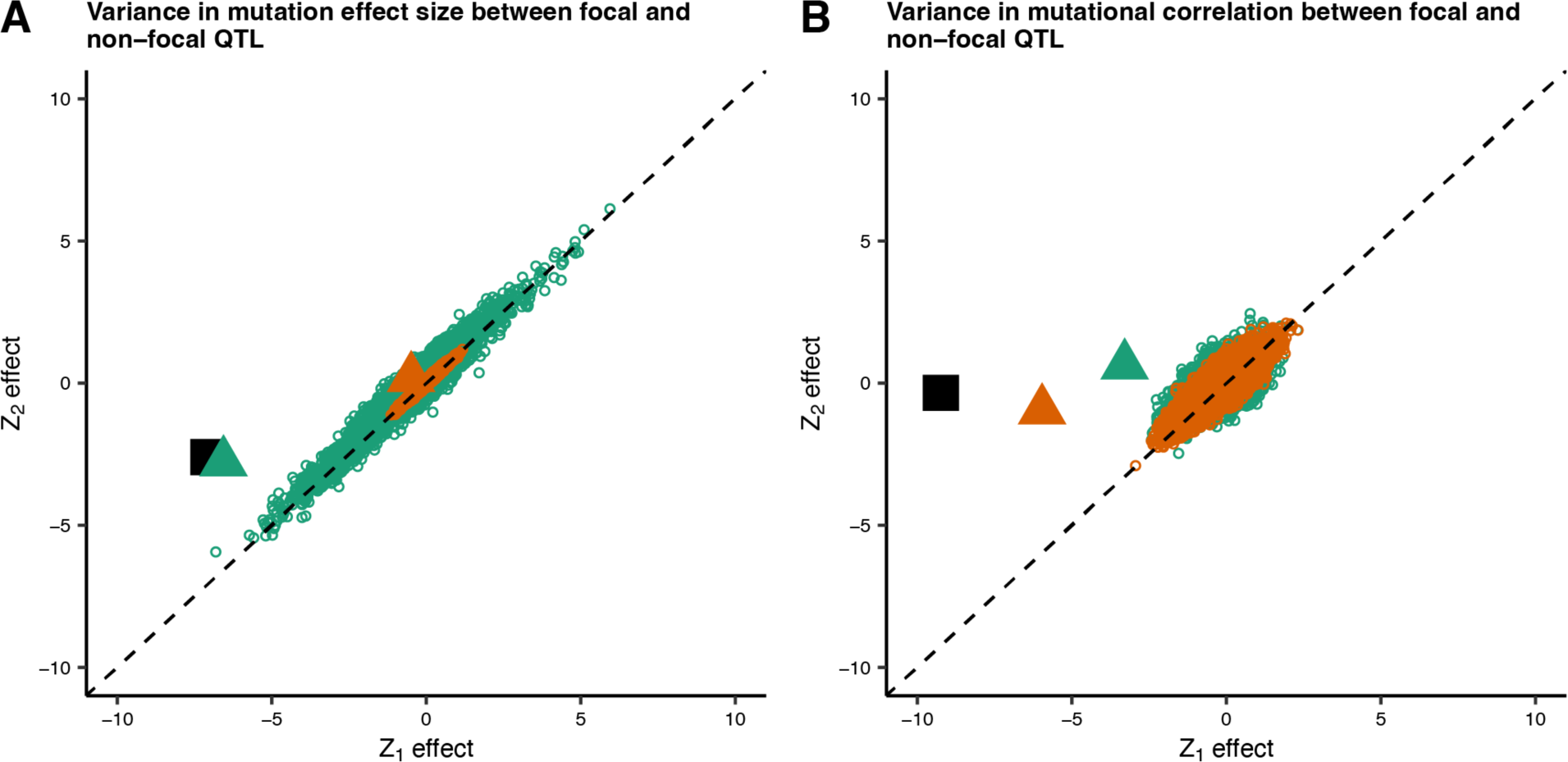
The mutations driving adaptation when one phenotype is under spatially divergent selection and the other is under spatially uniform selection (deleterious pleiotropy). Points show phenotypic effects for mutations segregating in *d_2_* across 1000 replicates for two example combinations of parameters (variance in mutation effect size, A: focal QTL *σ^2^* = 5; non-focal QTL *σ^2^*= 0.1; mutational correlations = 0.99 at all QTL; variance in mutational correlation, B: *σ^2^* = 0.5 for all QTL; focal QTL mutational correlations = 0.75; non-focal QTL mutational correlations = 0.9). Green points represent mutations at the focal QTL; orange points represent mutations at non-focal QTL. The mean divergence across replicates for focal and non-focal QTL is represented by green and orange triangles respectively, and the mean overall divergence by the black square. In both panels there are five QTL and migration rate = 0.005.

**Figure S3.**
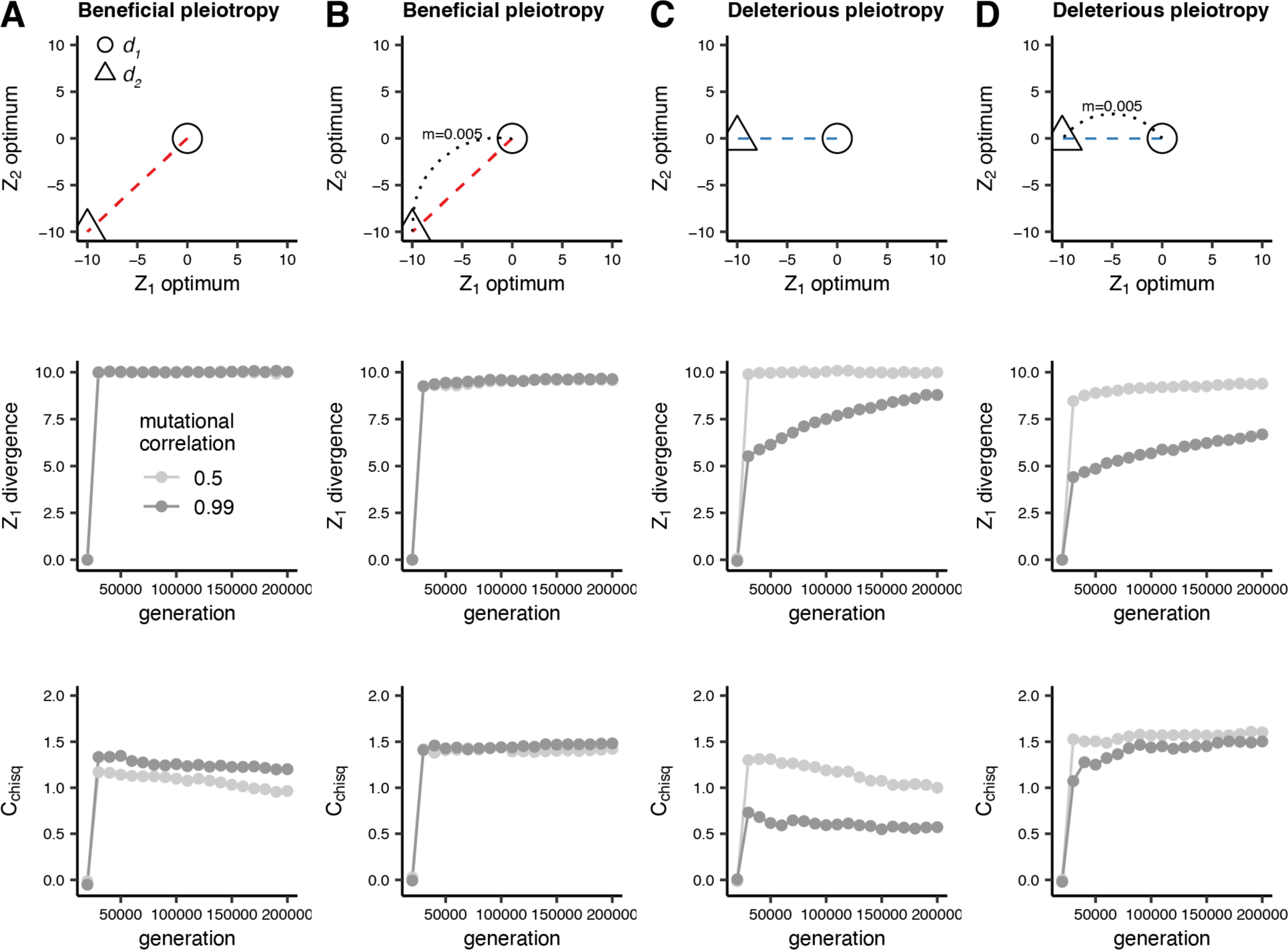
Divergence in *Z_1_* and repeatability (*C_chisq_*) over generations for four relationships between demes: where deme *d_2_* shifts to a new optimum in both traits *Z_1_* and *Z_2_* (divergent selection for both traits) without (A) and with (B) migration between *d_1_* and *d_2_*, and where *d_2_* shifts to new optimum in only trait *Z_1_* (divergent selection for one trait dimension and uniform selection for the other) without (C) and with (D) migration between *d_1_* and *d_2_*. *σ^2^* is 0.5 for the focal QTL and 0.1 for non-focal QTL. Mutational correlations are uniform at all QTL. These simulations use two phenotypes (one with divergent optima and the other with one non-divergent optima), and were run for 200,000 generations and 1000 replicates, with statistics calculated across replicates.

**Figure S4.**
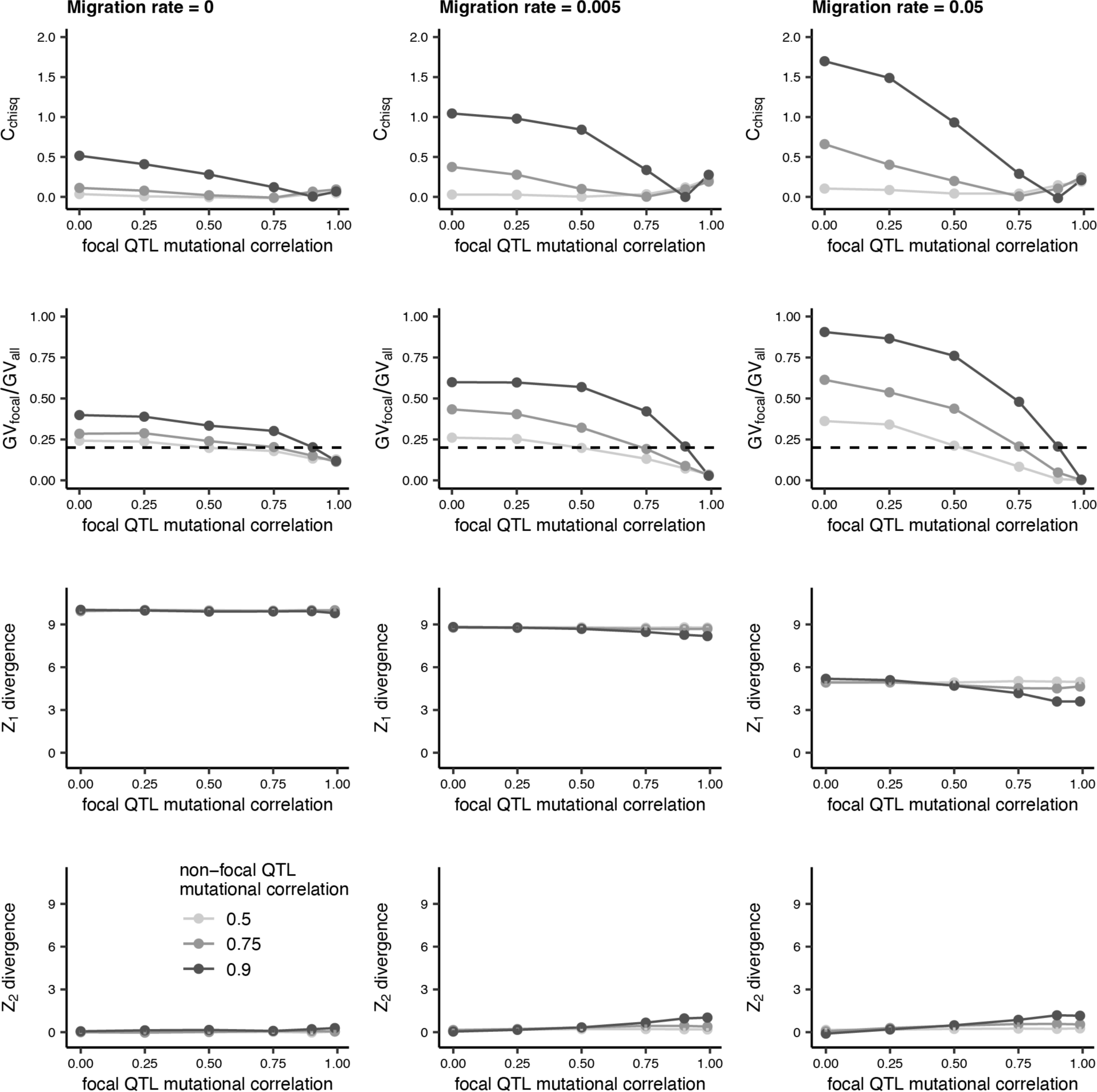
Repeatability (*C_chisq_*) in *Z_1_* against focal QTL mutational correlation for varying values of non-focal QTL mutation correlation (top row), the mean proportion of all *GV* explained by *GV* at the focal QTL (second row), and divergence between demes in *Z_1_* (third row) and *Z_2_* (bottom row). These simulations use deleterious pleiotropy an *σ^2^* of 0.1 and two phenotypes (one divergent and one non-divergent). They were run for 20,000 generations and 1000 replicates, with statistics calculated across replicates.

**Figure S5.**
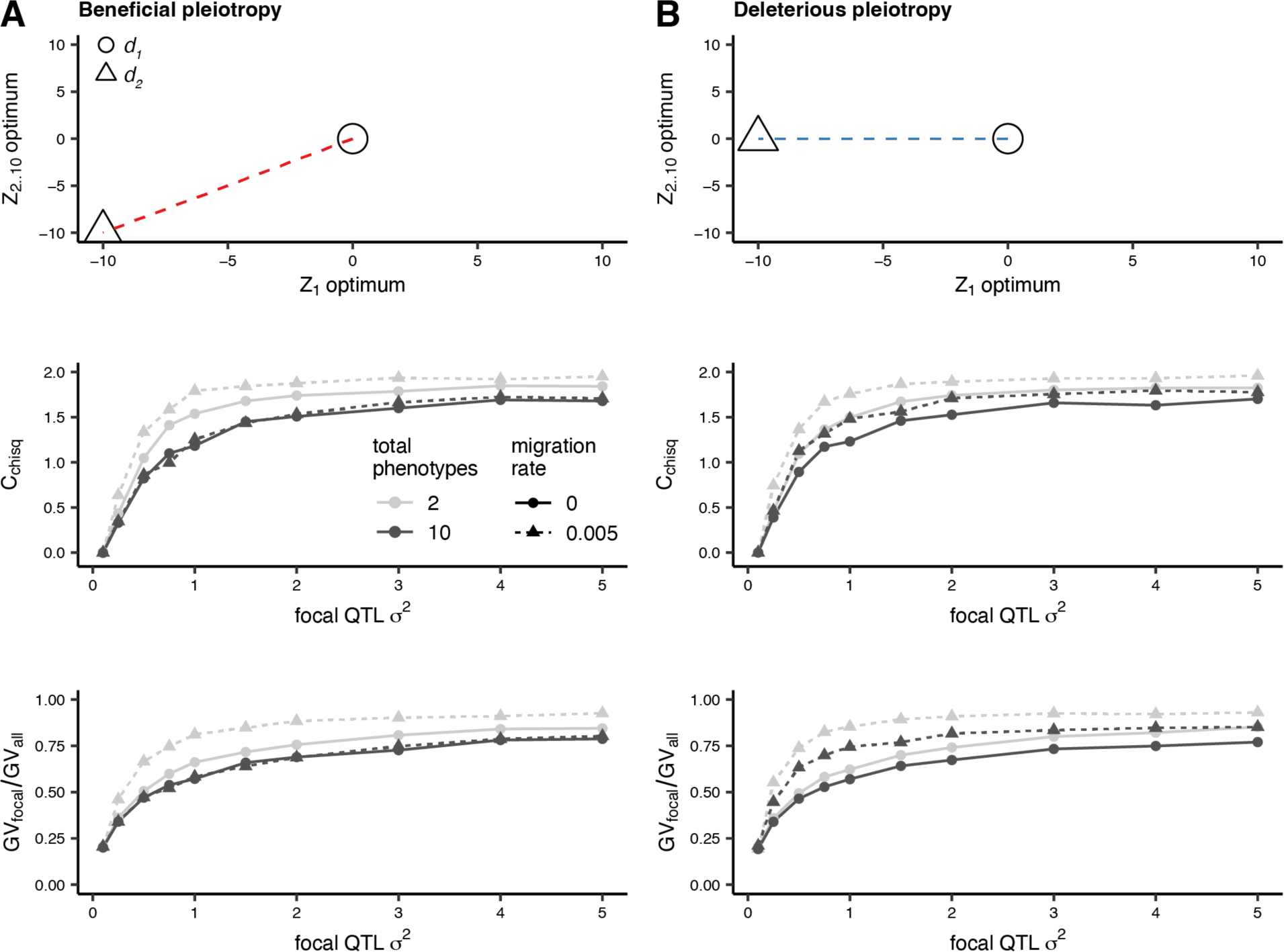
The effects of increasing trait dimensionality on repeatability (*C_chisq_*) arising from variation in *σ^2^*at the focal QTL where deme *d_2_* shifts to a new optimum in both trait dimensions *Z_1_* and *Z_2_* (divergent selection for both traits) with and without migration between *d_1_* and *d_2_* (A), and where *d_2_* shifts to new optimum in only *Z_1_* (divergent selection for one trait and uniform selection for the other) with and without migration between *d_1_* and *d_2_* (B). *σ^2^* at the focal QTL is varied, *σ^2^* for non-focal QTL is 0.1 and mutational correlations at all QTL are 0. Simulations were run for 20,000 generations and 1000 replicates, with statistics calculated across replicates.

**Figure S6.**
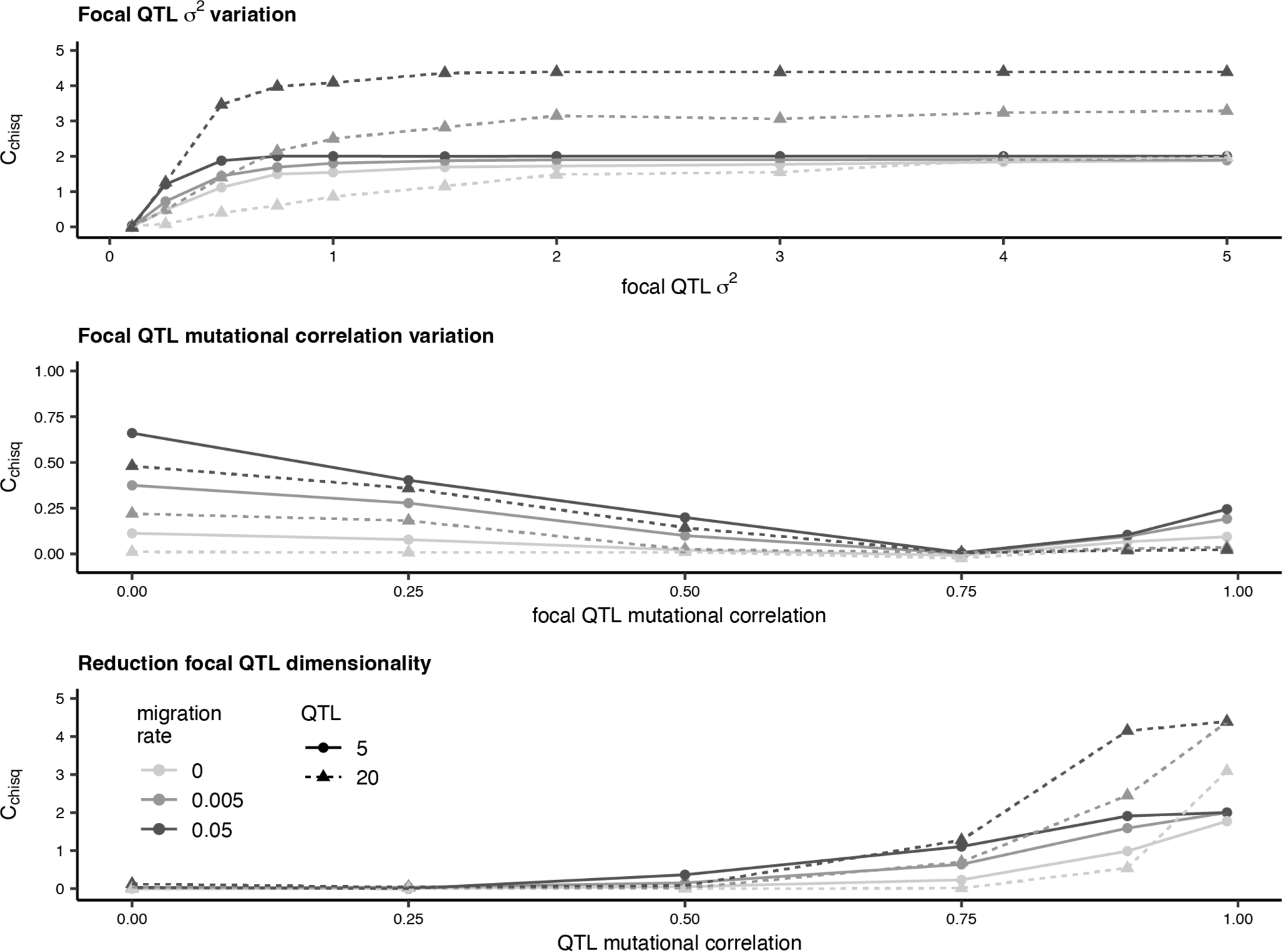
Effects of increasing the number of QTL modelled from five (solid lines, circle points) to 20 (dashed lines, triangle points). In the top pane we examine effect-magnitude variation at the focal QTL (as in Fig. 2), with mutational correlations for all QTL fixed at 0.5. In the middle pane we examine mutational correlation variation at the focal QTL (as in Fig. 3), with mutational correlations at non-focal QTL of 0.75 and *σ^2^* at 0.5. In the lower pane we examine a reduction in dimensionality at the focal QTL (as in Fig. 5), where the total number of phenotypes is two and *σ^2^* is 0.5. These simulations use deleterious pleiotropy and two phenotypes (one divergent and one non-divergent). They were run for 20,000 generations and 1000 replicates, with statistics calculated across replicates.

## Notes

https://github.com/pbattlay/pleio-sims/

### Competing Interest Statement

The authors have declared no competing interest.

### Summary of Updates

Substantial changes to text to increase clarity. Increased simulation replication.

